# Neuroendocrine negative SCLC is mostly RB1 WT and may be sensitive to CDK4/6 inhibition

**DOI:** 10.1101/516351

**Authors:** Dmitriy Sonkin, Suleyman Vural, Anish Thomas, Beverly A. Teicher

## Abstract

Small cell lung cancer (SCLC) is an aggressive form of lung cancer with limited therapeutic options, a very high mortality rate and is characterized, in most cases, by neuroendocrine features. A small but important subset of SCLC has intact RB1. However, other characteristics of this subset are not well-defined. To comprehensively assess the underlying genomics of SCLC cell lines with functional RB1, we examined 48 SCLC cell lines from the Cancer Cell Line Encyclopedia (CCLE) collection. Out of these 48 SCLC cell lines, 8 were found to be RB1 WT. We found that RB1 WT SCLC lines are enriched for loss of neuroendocrine lineage markers with CDKN2A inactivation, or CCND1 amplification. Six RB1 WT SCLC cell lines were included in NCI SCLC drug sensitivity screen and two of them were sensitive to CDK4/6 inhibition.

## Introduction

Small cell lung cancer (SCLC) is an aggressive form of lung cancer with limited therapeutic options, a very high mortality rate and is characterized, in most cases, by neuroendocrine features. The majority of SCLC are genetically characterized by homozygous inactivation of RB1 and TP53 tumor suppressor genes (1, 2, 3, 4, 5, 6). The prevailing hypothesis is that inactivation of RB1 in SCLC leads to increase in cellular proliferation due to loss of cell cycle control and inactivation of TP53 prevents oncogene induced senescence. SCLC diagnosis is commonly based on morphological features of biopsy or cytology samples. A panel of neuroendocrine markers (CHGA, NCAM1, SYP) may also be utilized (7, 8). In a noticeable fraction of cases, SCLC is present along with another lung cancer subtype(s) such as: large cell neuroendocrine carcinoma, large cell carcinoma, adenocarcinoma, squamous cell carcinoma (9). A small but important subset of SCLC has intact RB1 (10). However, genomic characteristics and potential therapeutic vulnerabilities of this subset are not well-defined.

## Materials and Methods

To comprehensively assess the underlying genomics of SCLC cell lines with functional RB1, we examined 52 SCLC cell lines from the Cancer Cell Line Encyclopedia (CCLE) collection (Supplemental Table S1) (11). These SCLC lines have comprehensive, high quality publicly available genetic and genomic data such as partial exome sequencing, whole exome sequencing, RNA-seq, AFFY SNP 6.0 arrays, and AFFY U133Plus2 mRNA arrays. NCI-H1339 cell line has been recently identified as potentially problematic due to stock contamination with other cell line(s) in some cell lines repositories (12), NCI-H1694 cell line has RB1 variant of uncertain significance, SHP-77 and DMS454 have low level of RB1 mRNA expression, to avoid potential confounding problems these 4 cell lines were excluded from this analysis.

## Results

### RB1 status

RB1 can be inactivated by multiple mechanisms, such as loss of function point mutations, deletions, insertions, exon inversions, splice site mutations, and loss of mRNA expression (due to promoter methylation or other mechanisms) (13, 14). All mechanisms of RB1 inactivation were taken in the account to determine the RB1 status. In addition to computational checks, RNA-seq data were manually reviewed to be certain, that there were no alterations of RB1 mRNA transcript. The importance of careful review of RB1 status is illustrated by the cell line COR-L24 which has a RB1 splice site mutation NM_000321.2:c.1389+5G>C. This splice site mutation is more than 3 bases away from an exon intron boundary, and therefore is not considered a canonical splice site mutation and may be missed by variant callers. This mutation leads to skipping of exon 14 as illustrated in Supplemental Figure S1. Out of 48 SCLC cell lines, 8 were found to be RB1 WT and are listed in Table 1. All 8 RB1 WT SCLC cell lines have RB1 protein expression (15, 16, 17, 18, 10, 19).

**Table 1.** RB1 WT SCLC cell lines

| Name | ASCL1 # | INSM1 # | NEUROD1 # | CHGA # | NCAM1 # | EGFR # | Key alterations |
| --- | --- | --- | --- | --- | --- | --- | --- |
| NCI-H2286 | 6 | 1 | 9 | 21 | 6 | 1139 | TP53 K291* hom, CDKN2A E33* hom |
| NCI-H841 | 24 | 3 | 12 | 21 | 8 | 1511 | TP53 C242S hom, SMARCA4 K934* hom |
| SW1271 | 14 | 3 | 1 | 8 | 15 | 1414 | TP53 C277F hom, NRAS Q61R, CDKN2A hom deletion |
| DMS-114<sup>^</sup> | 23 | 7 | 30 | 160 | 19 | 52 | TP53 R213* hom, SMARCA4 E1310* hom, CCND1 9 copies |
| SBC-5 | 13 | 24 | 12 | 34 | 15 | 38 | TP53 R248L hom, SMARCA4 AA1243fs hom |
| NCI-H1341 | 6 | 122 | 12 | 1554 | 408 | 2446 | PIK3CA E542K, MYC 5 copies |
| NCI-H211<sup>^</sup> | 25 | 127 | 6 | 29 | 1506 | 19 | TP53 R248Q hom, MYC 5 copies, CCND1 4 copies |
| DMS-53 | 15743 | 1464 | 375 | 357 | 310 | 99 | TP53 S241F hom, KRAS 8 copies high mRNA expression |

### Neuroendocrine status

Regulators of neuroendocrine differentiation such as ASCL1, INSM1, and NEUROD1 are highly expressed in SCLC (20, 21, 22). However It has been documented over the years that a small fraction of SCLC tumors do not express neuroendocrine lineage markers and transcription factors and is known as SCLC *variant* (23, 20, 24, 10). Table 1 lists 8 SCLC cell lines with WT RB1 in the ascending order of INSM1 mRNA expression. Five of the 8 RB1 WT SCLC cell lines had no noticeable mRNA expression of the neuroendocrine markers ASCL1, INSM1, NEUROD1, CHGA or NCAM1. Two of the 8 RB1 WT cell lines had no noticeable mRNA expression of ASCL1 and NEUROD1 and very low INSM1 mRNA expression, but with mRNA expression of CHGA or NCAM1. (AFFY probes set for SYP has a low dynamic range, for that reason SYP mRNA expression is not included in Table 1; however, SYP is included in Supplemental Table S1). In contrast, out of 40 SCLC cell lines with inactivating RB1 alterations or loss of expression (Supplemental Table S1), only 1 had no or very low mRNA expression of ASCL1, INSM1, NEUROD1, CHGA and NCAM1. Over all 62.5% (5 of 8) of RB1 WT SCLC lines were neuroendocrine lineage marker-negative compared with 2.5% (1 out of 40) of RB1 inactivated SCLC lines. Out of 6 neuroendocrine lineage marker-negative SCLC lines, 5 were RB1 WT. This observation is illustrated in Figure 1 and suggests that SCLC lines which do not express neuroendocrine lineage markers are enriched for intact RB1. A similar pattern was recently reported by McColl et al. (2017) (10).

**Figure 1.**
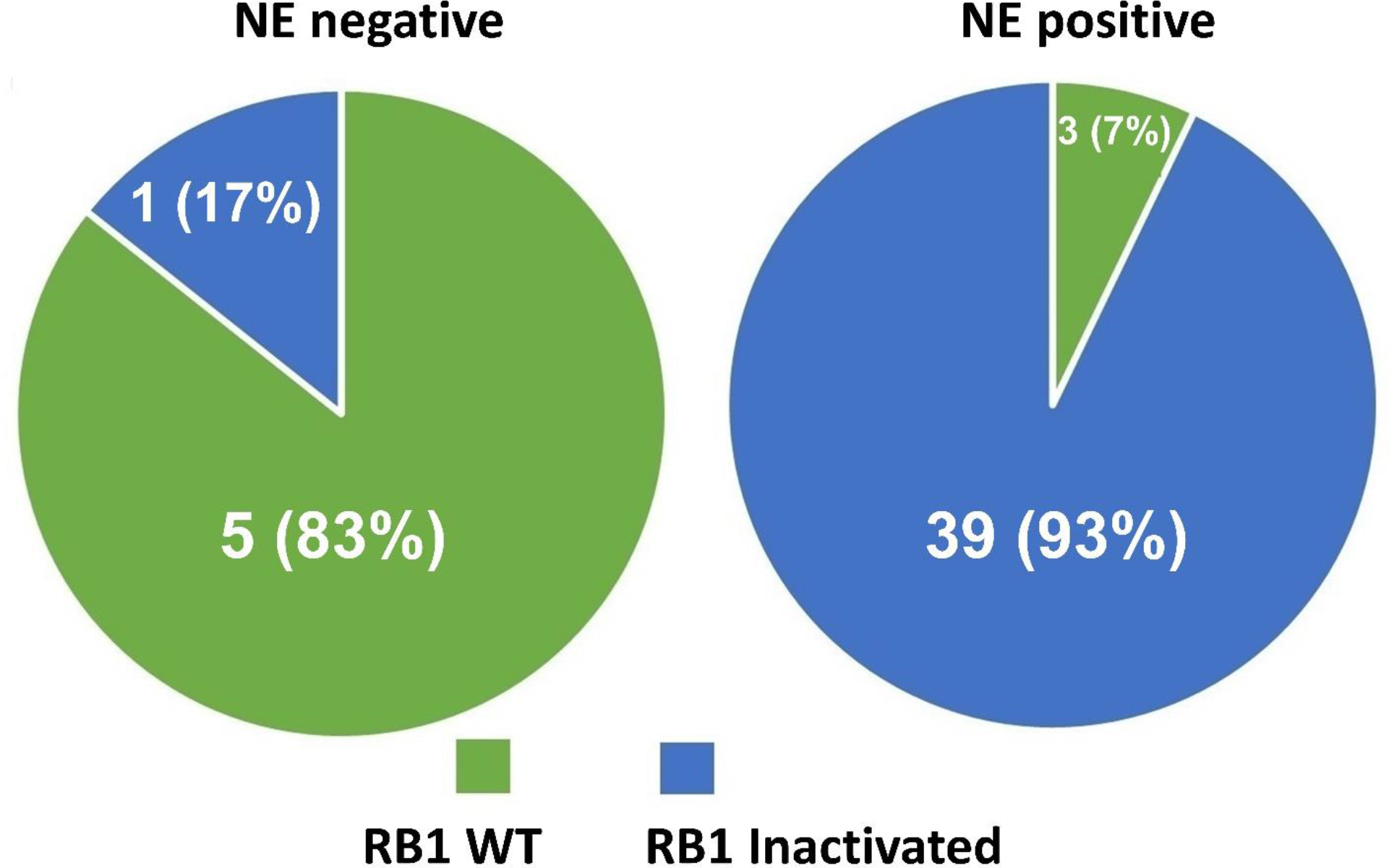
RB1 status of SCLC cell lines in relation to neuroendocrine (NE) status

### CDK4/6 inhibition

Out of the 48 CCLE SCLC cell lines, 35 were included in the NCI SCLC screen (25) of 420 approved and investigational oncology drugs. Due to their lack of functional RB1, the clear majority of SCLC cell lines were insensitive to the CDK4/6 inhibitor palbociclib (26, 27, 28). In contrast, 2 of 6 screened cell lines with functional RB1, DMS-114 and NCI-H211, were sensitive to the CDK4/6 inhibitor. Interestingly, these were the only SCLC cell lines with CCND1 amplification in set of 48 CCLE SCLC cell lines. Palbociclib concentration/response in the NCI SCLC screen is illustrated in Figure 2.

**Figure 2.**
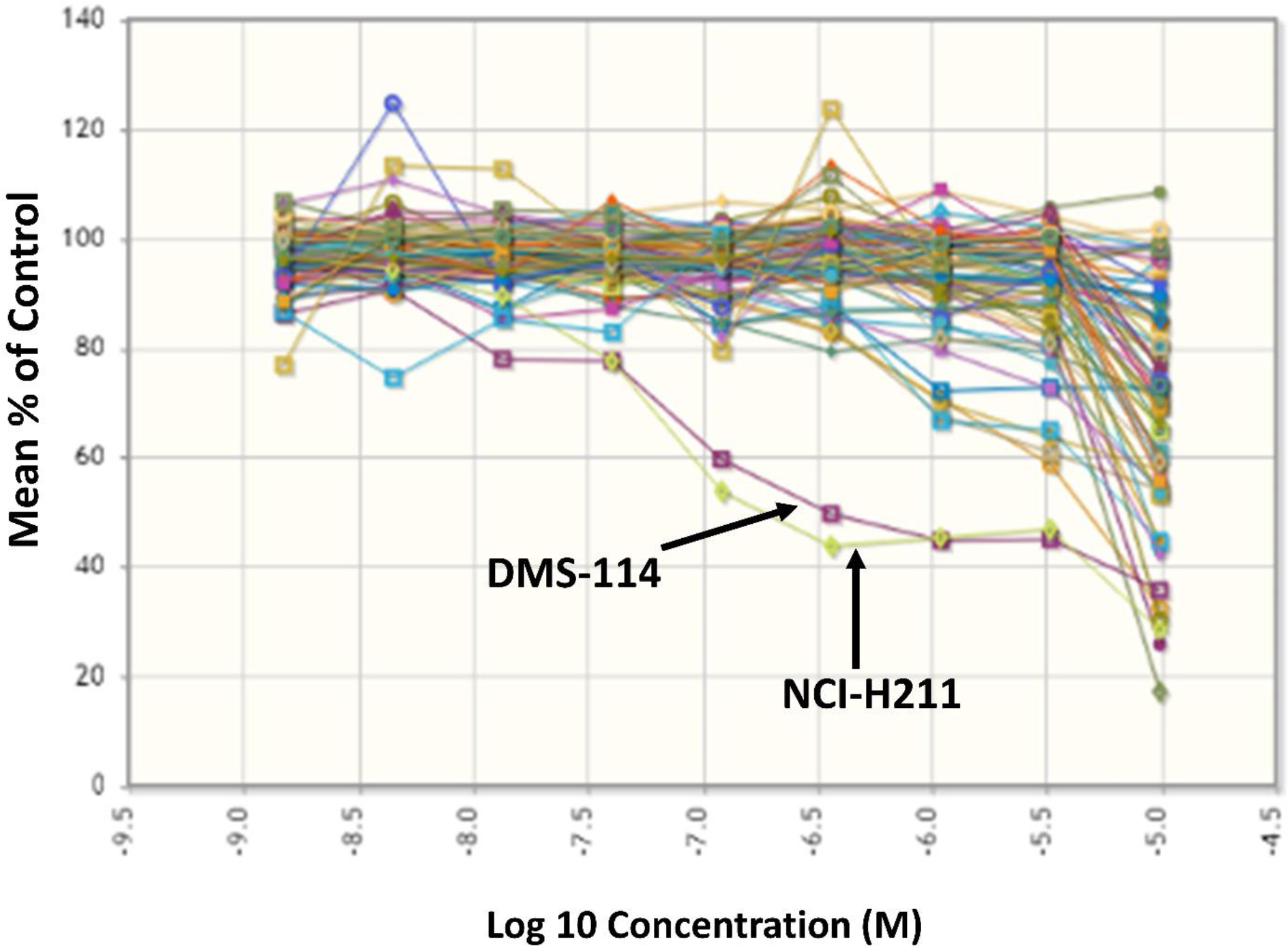
Palbociclib Concentration/Response in NCI SCLC screen

DMS-114 and NCI-H211 are also sensitive to CDK4/6 inhibitor palbociclib based on work by McColl et al. (2017) (10) and Barretina et al. (2012) (11). Palbociclib screening across CCLE (11) cell lines of different tumor types is illustrated in Figure 3. One of the 2 cell lines, DMS-114 has no or low mRNA expression of all five neuroendocrine lineage markers and the other NCI-H211 has no or low mRNA expression of 4 neuroendocrine lineage markers and has mRNA expression of NCAM1.

**Figure 3.**
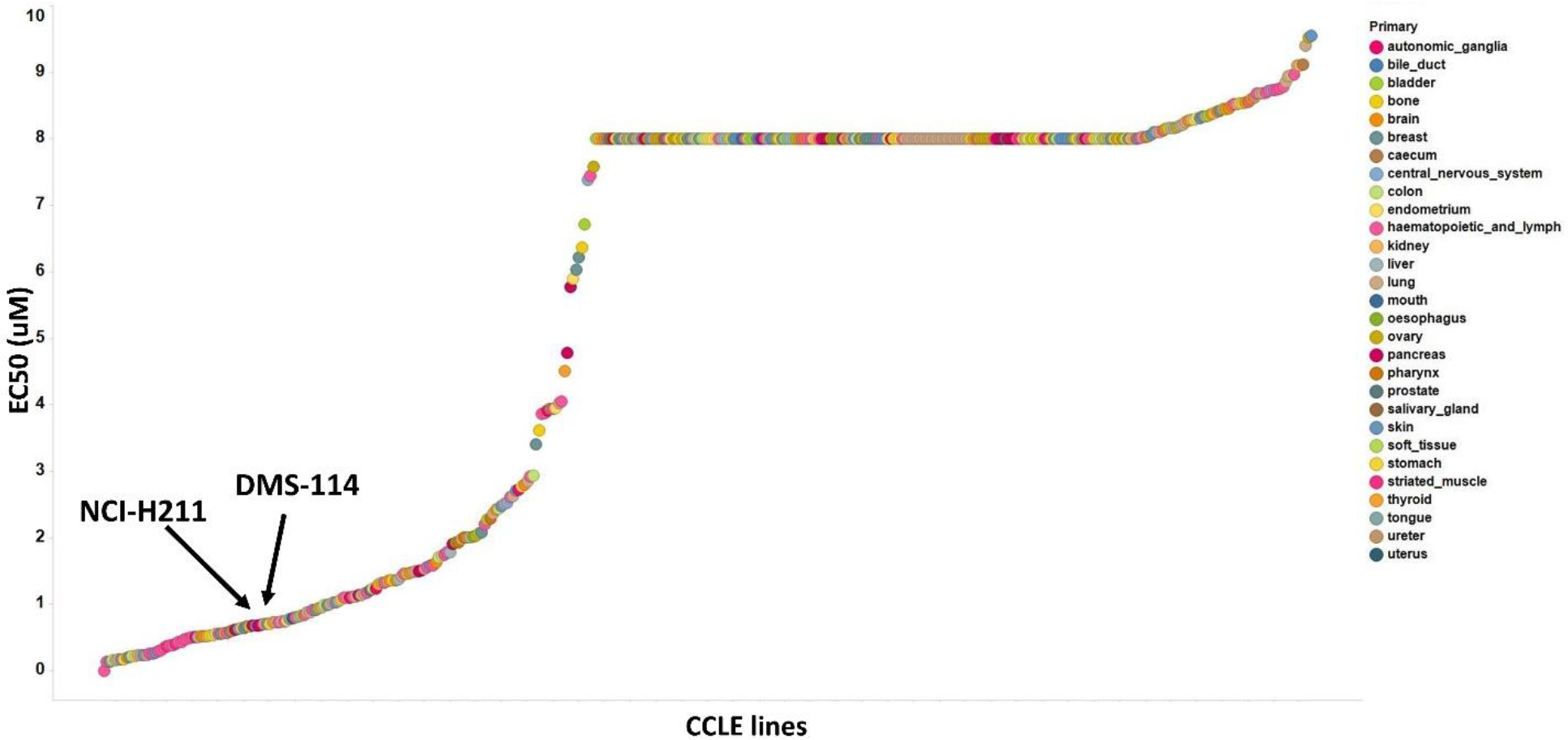
Palbociclib EC50 (μM) in CCLE screen

### Oncogenic alterations in WT RB1 SCLC

Oncogenic alterations in genes such as KRAS, NRAS, SMARCA4, CCND1, and CDKN2A are rare in SCLC; however, as can be seen from Table 1, these genes have oncogenic alterations in 7 of 8 RB1 WT SCLC cell lines. High levels of EGFR mRNA expression are rare in SCLC; however, as can be seen from Table 1, high EGFR gene expression is present in about half of the RB1 WT SCLC cell lines. It is interesting to note that these oncogenic alterations and high levels of EGFR mRNA expression frequently occur in non-small cell lung cancer (NSCLC) tumors (29). There are a few potential explanations for these observations. CCND1 amplification and CDKN2A inactivation may play similar role to RB1 inactivation resulting in cell cycle control defects. It is also possible that these alteration(s) originated in pulmonary neuroendocrine progenitor cells, but oncogene induced transformation and underlying lineage plasticity led to, at least, partial loss of the neuroendocrine phenotype. It is also possible that these alteration(s) originated in a common progenitor cell, which led to an unusual phenotype. It is also possible that some of SCLC lines may have a different cell of origin. For example, it was recently proposed that some of neuroendocrine marker negative SCLC may have originated from tuft cells (30). POU2F3, TRPM5, GNG13 are considered tuft cells markers, interestingly the NCI-H211 SCLC line, which is a typical neuroendocrine marker negative RB1 WT line, has comparatively high mRNA expression of these 3 markers. It is also potentially plausible that, in some of these cell lines, the original tumor had a mixed SCLC/NSCLC phenotype and after passaging in culture the NSCLC clones remained and the SCLC clones were lost.

## Discussion

### CCND1 amplification and CDKN2A inactivation

As mentioned above CCND1 amplification and CDKN2A inactivation may play similar role to RB1 inactivation resulting in cell cycle control defects, 4 out of 8 RB1 WT cell lines have CDKN2A inactivation or CCND1 amplification. In the prospective MSK-IMPACT study (31), out of 82 SCLC tumors, 16 did not have RB1 inactivation based on NGS DNA sequencing MSK-IMPACT-SCLC (It is possible that some of these tumors may have RB1 inactivated due to loss of mRNA expression or complex rearrangement). Two had CCND1 amplification and another 2 had homozygous CDKN2A deletion. In another study of 110 SCLC tumors, 10 had WT RB1 (George et al. (2015) Figure 1) and at least 2 of them had high CCND1 mRNA expression due to genetic alterations and both also had loss of CDKN2A mRNA expression (1). In the same study, out of 81 tumors with RNA-seq data, 3 had loss of neuroendocrine lineage markers (Supplemental Table S2), 2 of these tumors also had high CCND1 mRNA expression and 1 or 2 of these tumors may have had remaining functional RB1.

### Therapeutic implications

RB1 WT SCLC cell lines had noticeably lower levels of DLL3 mRNA expression than RB1 altered SCLC cell lines (Supplemental Table S1). A DLL3-targeted antibody-drug conjugate rovalpituzumab tesirine is being evaluated in a number of SCLC clinical trials. Previous reports have also shown neuroendocrine lineage marker negative SCLC cell lines to be associated with low DLL3 expression (31, 32). Therefore, RB1 WT SCLC cell lines are less likely to respond to DLL3-targeted drugs. However, this may present an opportunity to identify patients with WT RB1 among patients screened for DLL3 ADC clinical trials and having no or very low DLL3 expression.

Immune check point inhibitors are also currently undergoing clinical trial (NCT1928394, NCT02054806) in SCLC (34). Higher mutational load has been associated with greater efficacy of immune check point inhibitors (34, 35). The mutational load of RB WT SCLC cell lines did not to differ significantly from those of RB inactivated SCLC cell lines. The mutational load of SCLC cell lines NCI-H2286 and SW1271 is at the higher end of the spectrum; the mutational load of SCLC cell lines SBC-5 and NCI-H196 is at the middle of the spectrum and the mutational load for SCLC cell lines DMS114 and NCI-H841 is at the lower end of the spectrum (11).

## Conclusions

We draw attention to a subset of small cell lung carcinomas with loss of the typical neuroendocrine markers known as *variant* SCLC. Such tumors seem to be enriched for remaining functional RB1 alleles. This genomic subgroup of SCLC may be therapeutically relevant; RB1 WT SCLC are less likely to respond to DLL3 targeted drugs and may respond to CDK4/6 inhibitors. Based on McColl et al. (2017), a clinical trial of an CDK4/6 inhibitor in RB1 WT SCLC is being developed at Case Western Reserve University (personal communication).

## Supplemental Data

**Supplemental Table S1:** CCLE SCLC cell lines Genomic data

**Supplemental Table S2:** RNA-seq RPKM mRNA expression values for genes of interest and RB1 status from George et al., Nature, 2015

**Supplemental Figure S1:**
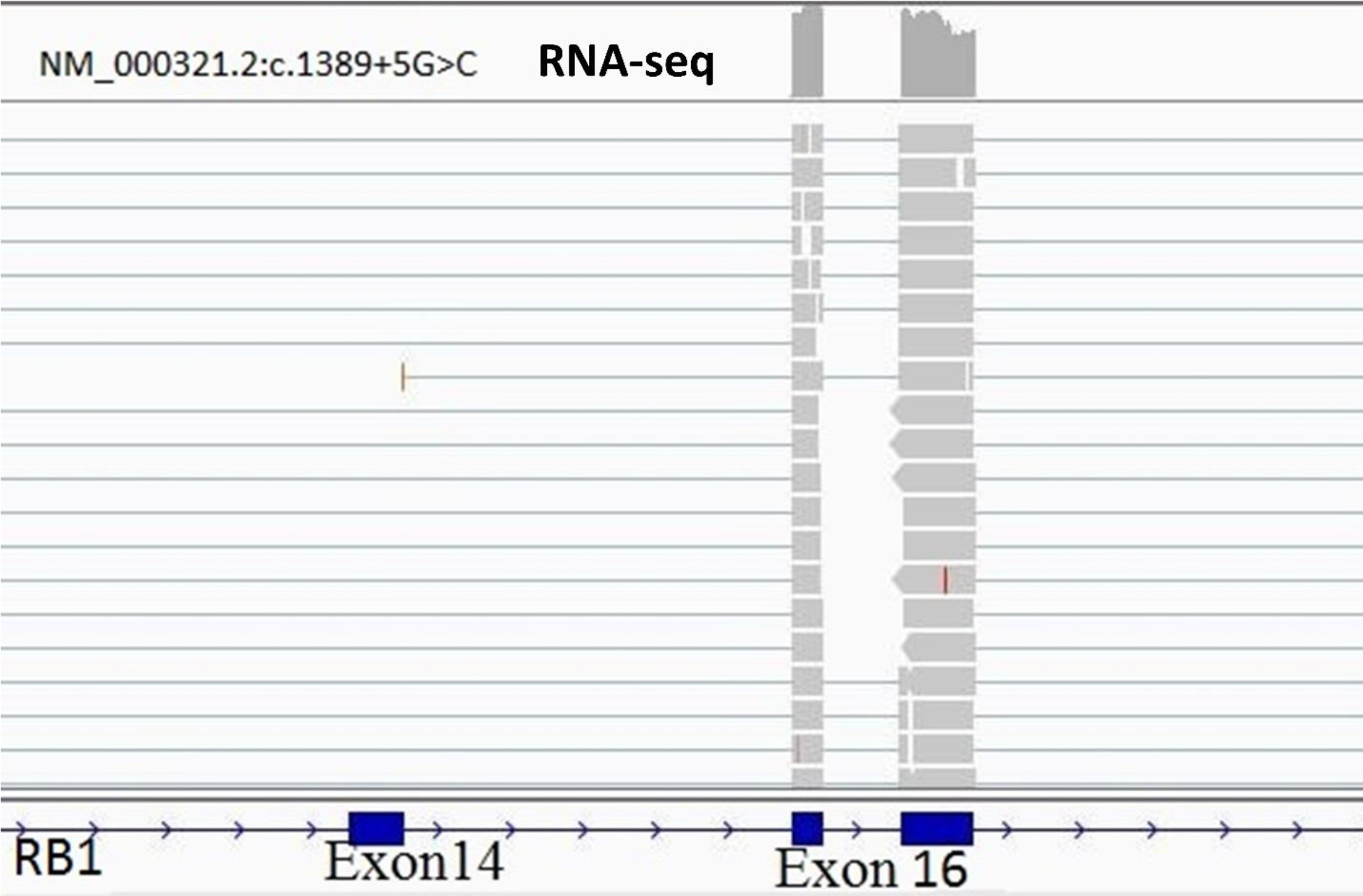
Example of splice site mutation >3 bases away from exon-intron boundary in COR-L24

